# A metagenomics-based tool for surveillance of bacterial pathogens in environmental samples

**DOI:** 10.64898/2025.12.16.694782

**Authors:** Chao Xiong, Min Gao, Brajesh K. Singh

**Author notes:** **Corresponding Author**: Brajesh K. Singh and Chao Xiong, |.

## Abstract

Bacterial pathogens are responsible for millions of human deaths and significant losses in food production annually. Soil acts as a natural reservoir for many critical bacterial pathogens, however, the lack of accurate, high-throughput methods to identify the distribution of bacterial pathogens in soil and other environments greatly limits our ability to develop effective surveillance and disease management strategies and policies. Here, we present MetaPatho, an easy-to-use, rapid, robust, and shotgun metagenomics-based tool for identifying human and plant bacterial pathogens in environmental metagenomes. We first constructed high-quality pathogenic databases for the identification of human and plant bacterial pathogens. In-silico tests and soil spike-in experiments, including pathogen DNA spike-in and cell spike-in tests, were then employed to examine the accuracy and robustness of our tool. The in-silico tests suggested that the tool accurately identified pathogenic sequences, achieving an average accuracy of 97.4%, with a specificity of 98.1% and a sensitivity of 97.2%. Further analyses from soil spike-in experiments showed that the tool accurately detected bacterial pathogens at different concentrations, achieving an average accuracy of 96.3%. Overall, our tools can provide a high-throughput and accurate surveillance of bacterial pathogens in environmental metagenomes, contributing to an effective risk management associated with human and plant diseases.

## Introduction

Soil and plant microbiomes are closely linked to agricultural productivity, plant health, food security, and environmental and human health^1–3^. Because pathogenic microbes within microbiomes pose significant threats to agricultural production and human health, understanding their distribution patterns and environmental drivers in soil-plant systems is considered a crucial component in achieving “One Health” framework^4–6^. Human bacterial pathogens (HBP) are a significant burden of public health and are associated with many important infectious diseases, such as tuberculosis, pneumonia, leprosy, cholera, and plague^7–9^. For example, *Mycobacterium tuberculosis* accounts for ∼10.6 M infections and ∼1.6 M deaths every year^8^, and *Mycobacterium leprae* is responsible for hundreds of thousands of human infections annually^10^. A recent study has estimated the global mortality associated with infectious diseases at 7.7 million deaths caused by 33 bacterial pathogens, including *Staphylococcus aureus*, *Escherichia coli*, *Streptococcus pneumoniae*, *Klebsiella pneumoniae*, and *Pseudomonas aeruginosa*^7^.

In addition to human pathogens, crop diseases caused by plant bacterial pathogens (PBP) can also indirectly influence human health by affecting global agricultural production and food security^11–16^. Some critical plant diseases lead to substantial losses of crop health and can affect ecosystem productivity and socio-economic development, including bacterial wilt, potato scab, and crown gall tumor^11–17^. Soil serves as a natural reservoir and habitat for many bacterial pathogens^5,6,18,19^. Among these human and plant pathogens, many are capable of surviving and proliferating in soil, such as *Ralstonia solanacearum*, *Streptomyces scabies*, *Pseudomonas aeruginosa*, and *Burkholderia pseudomallei*^5,18,20,21^. Others may enter soils through various other pathways, such as human wastes and agricultural practices, or via atmospheric deposition, becoming potential sources of infection, including *Salmonella enterica* and *Escherichia coli*^5,22–25^. Consequently, the precise identification and monitoring of bacterial pathogens in the soil and other environments are of critical importance for global public health, sustainable agricultural production and primary productivity of natural ecosystems.

Existing laboratory-based pathogen surveillance necessitates the isolation of specific pathogens from environmental or clinical samples before further identification and characterization can be performed through molecular assays^26^. This process is time-consuming, resource-intensive and suffers from low efficiency in detection^26^. Similarly, identifying potential pathogens in environmental samples based on 16S rRNA metabarcoding lacks efficacy and specificity^26–28^. The increasing use of shotgun metagenomic next-generation sequencing (mNGS) technologies offers a promising opportunity for the high-throughput and accurate surveillance of environmental pathogens^26,28–30^. Concurrently, a variety of software and pipelines based on distinct algorithms have been developed for taxonomic annotation of metagenomic data, such as Kraken 2^31^, MetaPhlAn^32^, and MG-RAST^33^. However, there’s limited evidence on whether these tools, designed for taxonomic annotation across all microbes, can accurately detect pathogens in the complex environmental samples^26,34^. In addition, cloud-deployable platforms like IDseq^35^ and SURPI^36^ face limitations in processing a large volume of samples and providing detailed functional profiles. For example, when assessing soil health and evaluating the functional potential of microbiomes, it is essential to obtain not only information on pathogen distribution but also on the key functional traits of the microbiome (e.g., genes that promote or suppress diseases and those involved in elemental cycling and secondary metabolite synthesis). However, such information is often highly limited in existing pipelines and online platforms. Given that environmental, particularly soil, samples are very complex and diverse, constructing a reliable and high-throughput pipeline for identifying environmental bacterial pathogens represents an important challenge. Therefore, integrating a simple, user-friendly, and accurate pathogen annotation module into routine metagenomic functional analysis pipelines is needed to advance the discipline.

By integrating new high-quality bacterial pathogen genome databases and the Virulence Factor Database (VFDB)^37^ and the Pathogen-Host Interactions database (PHI-base)^38^ databases we developed a shotgun metagenomics-based pipeline (Fig. 1), the MetaPatho, which allows for the accurate identification of both human and plant bacterial pathogens and associated virulence factors simultaneously in environmental metagenomes. Our pipeline exhibits the following features and advantages: (1) the analysis is rapid and straightforward, allowing the pathogen annotation module and virulence factor annotation module to be conveniently integrated into standard metagenomic functional analysis processes; (2) establishment of a list and high-quality pathogenic databases for soil-inhabiting human (HBPDB) and plant (PBPDB) bacterial pathogens; (3) accuracy validation of our tool in detecting bacterial pathogens through both in-silico tests and soil spike-in experiments including pathogen DNA spike-in and cell spike-in tests. These results suggest that our metagenomics-based pathogen identification pipeline, MetaPatho (https://github.com/ChaoXiong-2024/MetaPatho), can provide an important tool to inform identity and distribution of bacterial pathogens and contribute to future pathogen surveillance for environmental and public health management and policy formulation.

**Fig. 1.**
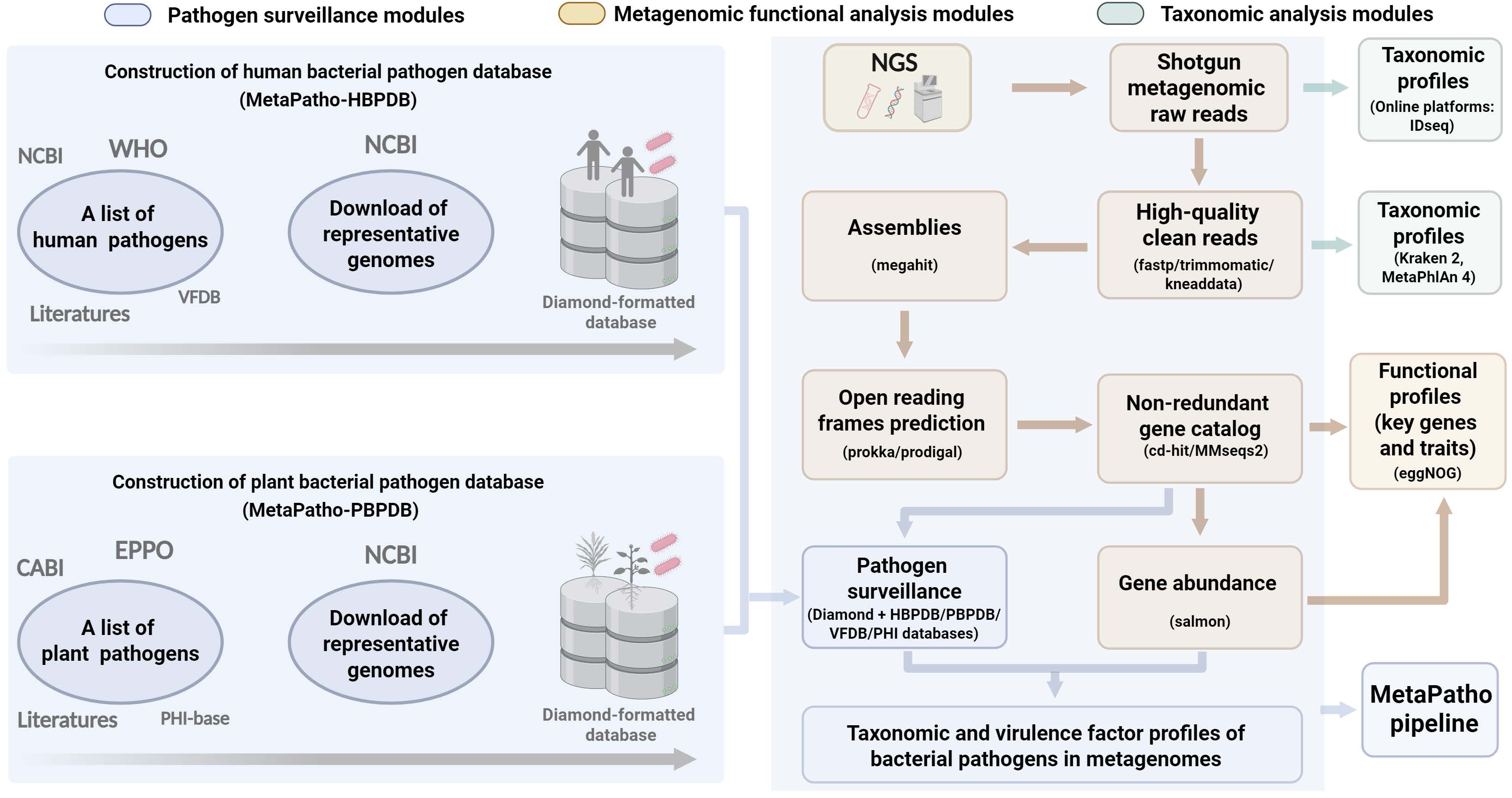
Construction of pathogenic databases and bioinformatic pipeline for pathogen surveillance. To ensure that pathogen identification pipeline remains accurate while being simple and user-friendly, we developed a high-quality bacterial pathogen genome database and easily integrate the pathogen annotation module into routine metagenomic functional analysis workflows. Our metagenomics-based pipeline (i.e., MetaPatho) for pathogen surveillance primarily consists of two components. Firstly, we construct high-quality pathogenic databases for the identification of human and plant bacterial pathogens. Secondly, these databases were employed to identify pathogen sequences within metagenomic functional analysis pipelines. Compared with other available tools (e.g., Kraken 2 and MetaPhlAn 4), our pathogen identification pipeline is fully integrated into standard metagenomic analysis modules and does not require additional software installation. For example, when analyzing large sample batches, our pipeline only requires an additional annotation step against the pathogen database on top of standard metagenomic analyses, enabling simultaneous acquisition of pathogen distribution and microbiome functional profiles, such as genes involved in pathogen promotion and suppressions or involved with soil nutrient cycling.

## Results and discussion

### Construction of pathogenic databases

To ensure that pathogen identification pipeline remains accurate while being simple and user-friendly (e.g., avoiding the installation of complex dependencies), we developed a high-quality bacterial pathogen genome database and easily integrate the pathogen annotation module into routine metagenomic functional analysis workflows. This approach allows user to obtain pathogen distribution information simply by annotating against the pathogen database during standard metagenomic functional profiling, thus avoiding the need for new software installation and additional environment dependencies. Further to systematically identify and explore the distribution patterns of soil-inhabiting bacterial pathogens, we first targeted key bacterial pathogens that inhabit the soil as well as significant pathogens that may be transferred to soil through various biotic and abiotic pathways. First, we compiled a list of human bacterial pathogens using multiple authoritative sources, including the World Health Organization (WHO)’s recommended list^39,40^, the National Center for Biotechnology Information (NCBI) pathogen detection system (https://www.ncbi.nlm.nih.gov/pathogens/), VFDB^37^, and previous literature^4,5,18,19,41–43^ (Fig. 1). This integrative methodology ensured comprehensive coverage of key bacterial pathogens with direct implications for human health. Ultimately, our pathogen catalogue includes a total of 82 pathogen species across 44 genera and 33 families, including *Enterobacteriaceae*, *Mycobacteriaceae*, *Clostridiaceae*, *Vibrionaceae*, and *Staphylococcaceae* (Extended Data Table 1). We next downloaded high-quality representative genomes of the 82 pathogen species from the NCBI genome repository. In the case of pathogens frequently implicated in foodborne illnesses (such as *Salmonella enterica*, *Escherichia coli*, and *Listeria monocytogenes*) and leading pathogens (like *Pseudomonas aeruginosa*, *Staphylococcus aureus*, and *Burkholderia pseudomallei*), our strategy extended beyond just representative genomes. For these, we also sourced genomes from various strains and newly isolated environmental samples, aiming for a comprehensive screening. We then translated the nucleotide sequences of these pathogen genomes into protein sequences and constructed a reference pathogenic database for human bacterial pathogens (MetaPatho-HBPDB) using DIAMOND (Fig. 1)^44^.

Given that a majority of plant pathogens are soilborne, including *Ralstonia solanacearum*, *Streptomyces scabies*, and *Agrobacterium tumefaciens*, the high-throughput surveillance of these pathogens from soils emerges as an imperative task, particularly in the context of global changes that may exacerbate their prevalence and distribution^13,14,16^. Similar to what we did for human pathogens, we initially constructed a catalogue of significant plant bacterial pathogens based on the global database from European and Mediterranean Plant Protection Organization (EPPO)^45^, CABI^45^, PHI^38^, as well as previous literature^17,46,47^ (Fig. 1). This catalogue includes 113 bacterial pathogenic species from 27 genera across 14 families (Extended Data Table 2).

Subsequently, we downloaded representative genomes of these pathogens from the NCBI Genomic Database. For frequently reported significant plant pathogens, such as *Pseudomonas syringae*, *Ralstonia solanacearum*, *Agrobacterium tumefaciens*, *Xanthomonas oryzae* pv. *Oryzae*, *Xanthomonas campestris*, genomes of newly isolated environmental strains were also downloaded. For pathogens with multiple subspecies causing diseases in different crops, such as *Candidatus Liberibacter*, *Clavibacter michiganensis*, and *Pseudomonas syringae*, genomes of these various subspecies or pathovars were also downloaded. We next converted the nucleotide sequences of these genomes into protein sequences, and then employed DIAMOND to construct a reference pathogenic database for plant bacterial pathogens (MetaPatho-PBPDB) (Fig. 1).

### In-silico tests of the pathogen identification pipeline

In-silico tests were employed to evaluate the performance of the MetaPatho pipeline. We first constructed simulated datasets (i.e., MOCK community) for both the HBPDB and PBPDB by selecting randomly translated protein sequences of varying lengths from the genomes of 40 bacterial pathogens (serving as true positives) and 10 non-pathogenic bacteria (serving as true negatives) (Fig. 2). To enhance the representativeness of our simulated datasets, we included significant soil-borne pathogens (e.g., *Ralstonia solanacearum*, *Streptomyces scabies*, *Burkholderia pseudomallei*, and *Pseudomonas aeruginosa*) and those have a critical impact on human health (e.g., *Mycobacterium tuberculosis*) or frequently reported as foodborne pathogens (e.g., *Salmonella enterica*)^5,7–9,18,20^. Additionally, we selected pathogens with varying genome sizes, ranging from 1.1 (*Chlamydia psittaci*) to 8.4 (*Burkholderia cepacia*) Mb for HBPDB and from 1.1 (*Candidatus Liberibacter americanus*) to 11.2 (*Streptomyces acidiscabies*) Mb for PBPDB. We then identified pathogen sequences within the MOCK community using our pipeline. The results demonstrated that our pipeline accurately recognized pathogenic sequences of different lengths, achieving an average accuracy of 97.4% (using identify 90, human pathogen 99.3%, plant pathogen 95.4%), with a specificity of 98.1% (human pathogen 98.0%, plant pathogen 98.2%) and a sensitivity of 97.2% (human pathogen 99.7%, plant pathogen 94.7%) (Fig. 3). Additionally, we tested the accuracy and sensitivity of our pipeline at various identity (ID) thresholds (ID 70, ID 80, ID 90, and ID 99). The analysis results indicate that our pipeline performed well across various ID thresholds, achieving accuracies of 93.8-99.3%, specificities of 89.0-99.9%, and sensitivities of 94.7-99.7% (Fig. 3). The positive predicted value of our pipeline ranged from 97.2-99.9%, and the negative predict value was 80.1-98.9% (Fig. 3). All these results demonstrate that our pipeline can accurately identify pathogenic sequences within the MOCK community.

**Fig. 2.**
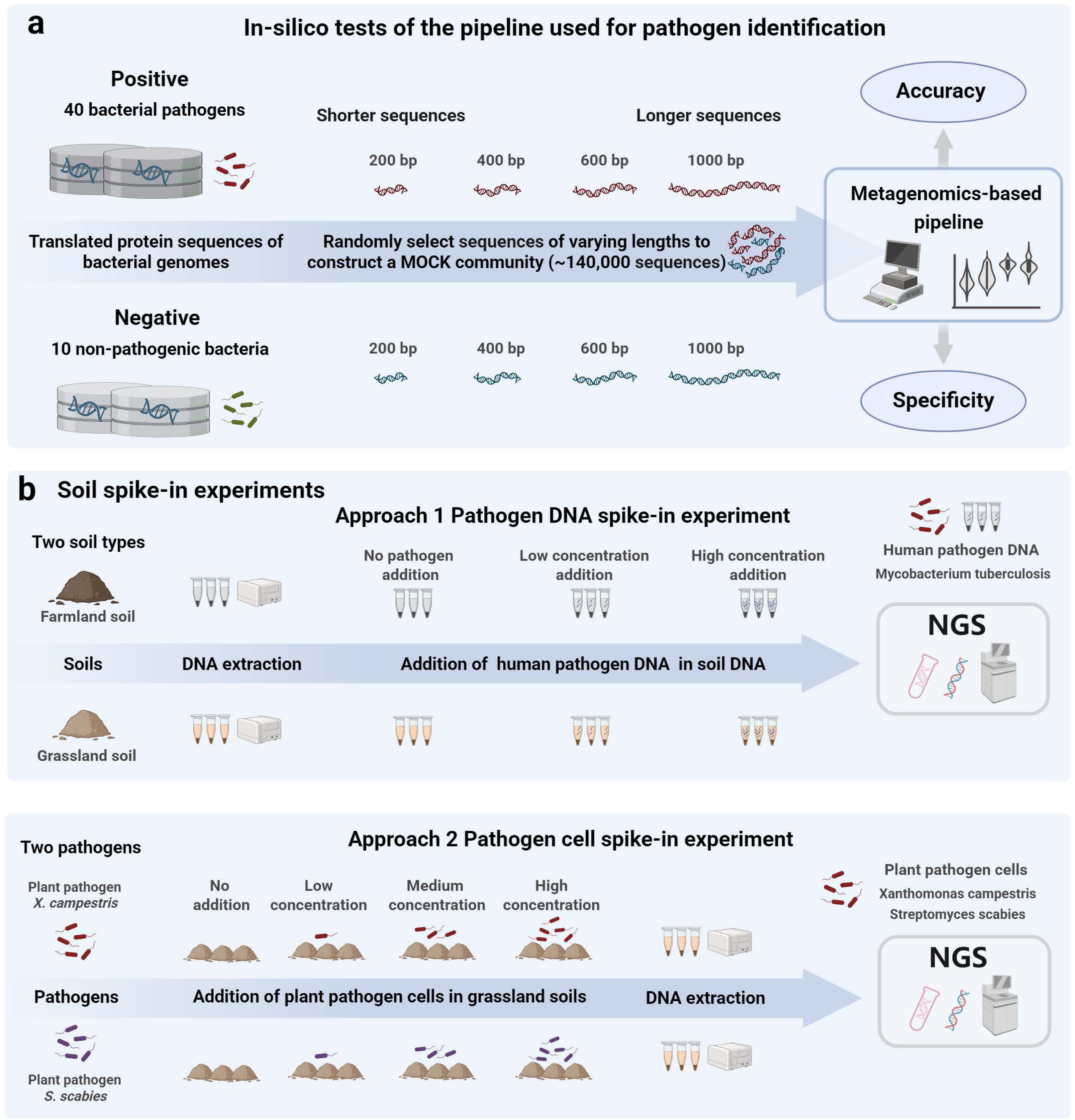
Validation of the accuracy of MetaPatho pipeline through in-silico tests and soil spike-in experiments. **a.** In-silico tests based on simulated dataset (i.e., MOCK community) were employed to assess the accuracy, specificity, and sensitivity of our pipeline. The resultant MOCK community comprised approximately 140,000 sequences, including around 100,000 sequences of 200 bp, 25,000 sequences of 400 bp, 10,000 sequences of 600 bp, and 500 sequences of 1000 bp. **b.** Soil spike-in experiments, including both pathogen DNA spike-in (approach 1) and cell spike-in tests (approach 2), were employed to provide evidence supporting the robustness of our pipeline.

**Fig. 3.**
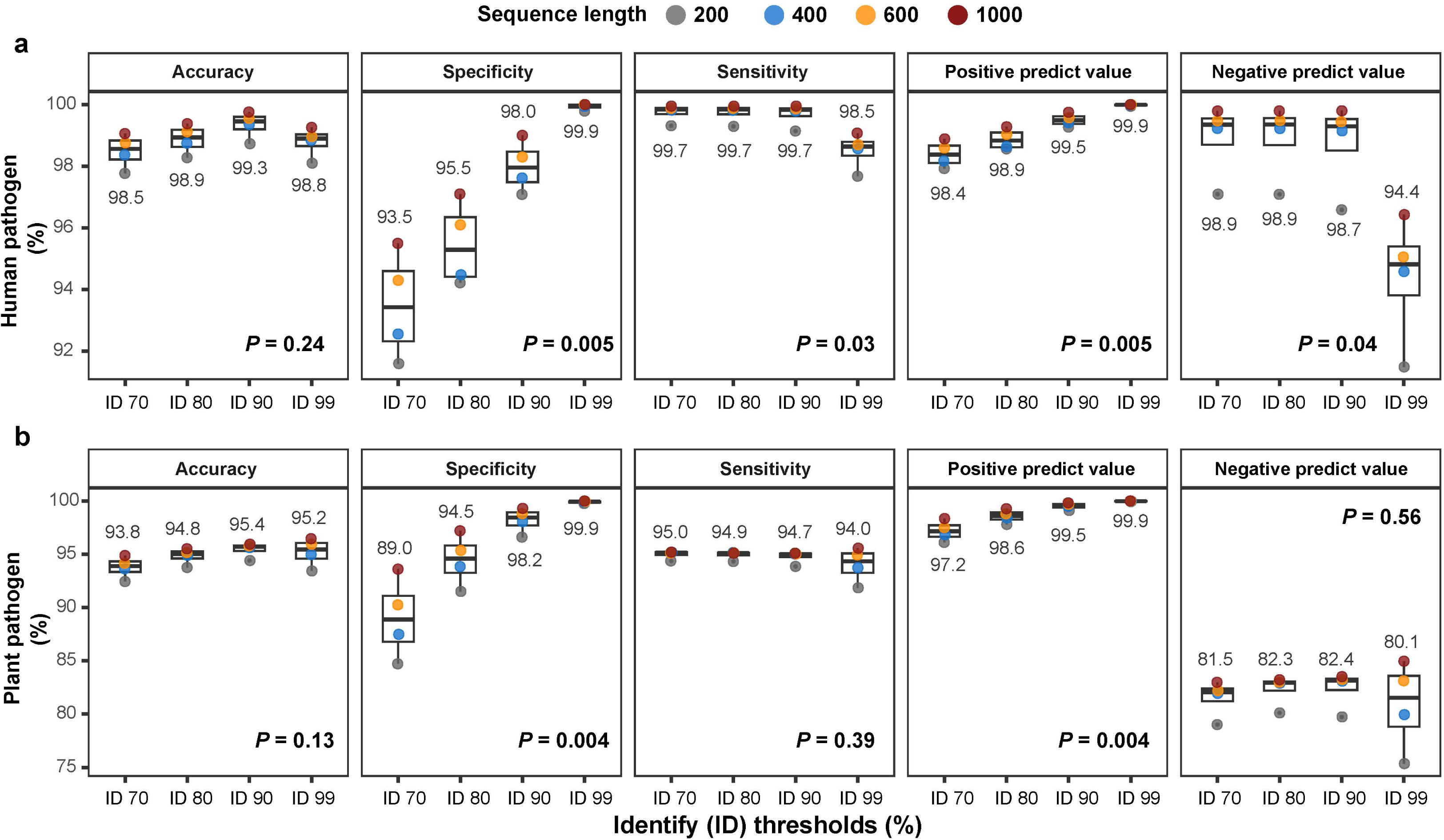
Performance of MetaPatho pipeline in in-silico tests. a-b. Box plots illustrating the performance of our pipeline in in-silico tests, encompassing accuracy, specificity, sensitivity (also known as recall), positive predictive value, and negative predictive value. We assessed the performance of our pipeline at various identify thresholds (ID 70, ID 80, ID 90, and ID 99). In box plots, the edges of the box show lower and upper quartiles, the middle line indicates the median. Accuracy, ACC, measures the overall correctness of the pipeline; Specificity, SP, measures the proportion of actual negatives that are correctly identified; Sensitivity, SE, measures the proportion of actual positives that are correctly identified; Positive predictive value, PPV, measures the proportion of positive identifications that are actually correct; Negative predictive value, NPV, measures the proportion of negative identifications that are actually correct. TP, true positive number; TN, true negative number; FP, false positive number; FN, false negative number. ACC = (TP+TN) / (TP+TN+FP+FN); SP = TN / (TN+FP); SE = TP / (TP+FN); PPV = TP / (TP+FP); NPV = TN / (TN+FN).

### Performance validation using pathogen DNA spike-in experiments

We then employed pathogen DNA spike-in experiments to provide evidence for the robustness of our pipeline. In this validation section, we selected key human bacterial pathogens *Mycobacterium tuberculosis* for experimental testing^8,40^. We opted to directly add purchased genomic DNA of human pathogens (*Mycobacterium tuberculosis* H37Ra, ATCC 25177) in soil DNA samples. Results of metagenomic analysis indicated that our pipeline can accurately detect pathogen DNA at different concentrations from two soil types (Fig. 4, Table 1). We calculated the accuracy of the pipeline using the defined formula (Fig. 4), and the results demonstrated that our pipeline achieves an average accuracy of 98.1% (using ID 90) in pathogen identification across both soil types (100% in farmland soil and 96.3% in grassland soil) (Fig. 4, Table 1). In farmland soil, the average relative abundance values (Reads Per Million, RPM) identified for pathogens were 25.2 (no pathogen addition), 33.8 (low concentration addition), and 152.0 (high concentration addition). For grassland soil, the average RPM values identified for pathogens were 48.7 (no pathogen addition), 59.4 (low concentration addition), and 87.0 (high concentration addition) (Fig. 4). We also explored the impact of employing different identity (ID) thresholds on the performance of our pipeline during sequence annotation with DIAMOND. Our results showed that at a higher ID threshold (99%), our pipeline achieved an accuracy of up to 100% (Extended Data Fig. 1, Table 1). When using a moderate ID threshold (80%), the pipeline maintained a high accuracy of 96.3% (Extended Data Fig. 1, Table 1). Together, these findings provide strong evidence that our tool can accurately identify human pathogens in soil. However, when using lower ID thresholds (60% and 70%), the average accuracy of our pipeline decreased to 74-87% (Table 1), highlighting the importance of selecting an appropriate ID threshold. Considering the high diversity and variability of soil microbiomes in real-world^5,48,49^, large-scale surveys, adopting a moderate ID threshold (i.e., 80-90%) could enable a more comprehensive and accurate characterization of potential human pathogen distributions in soil.

**Fig. 4.**
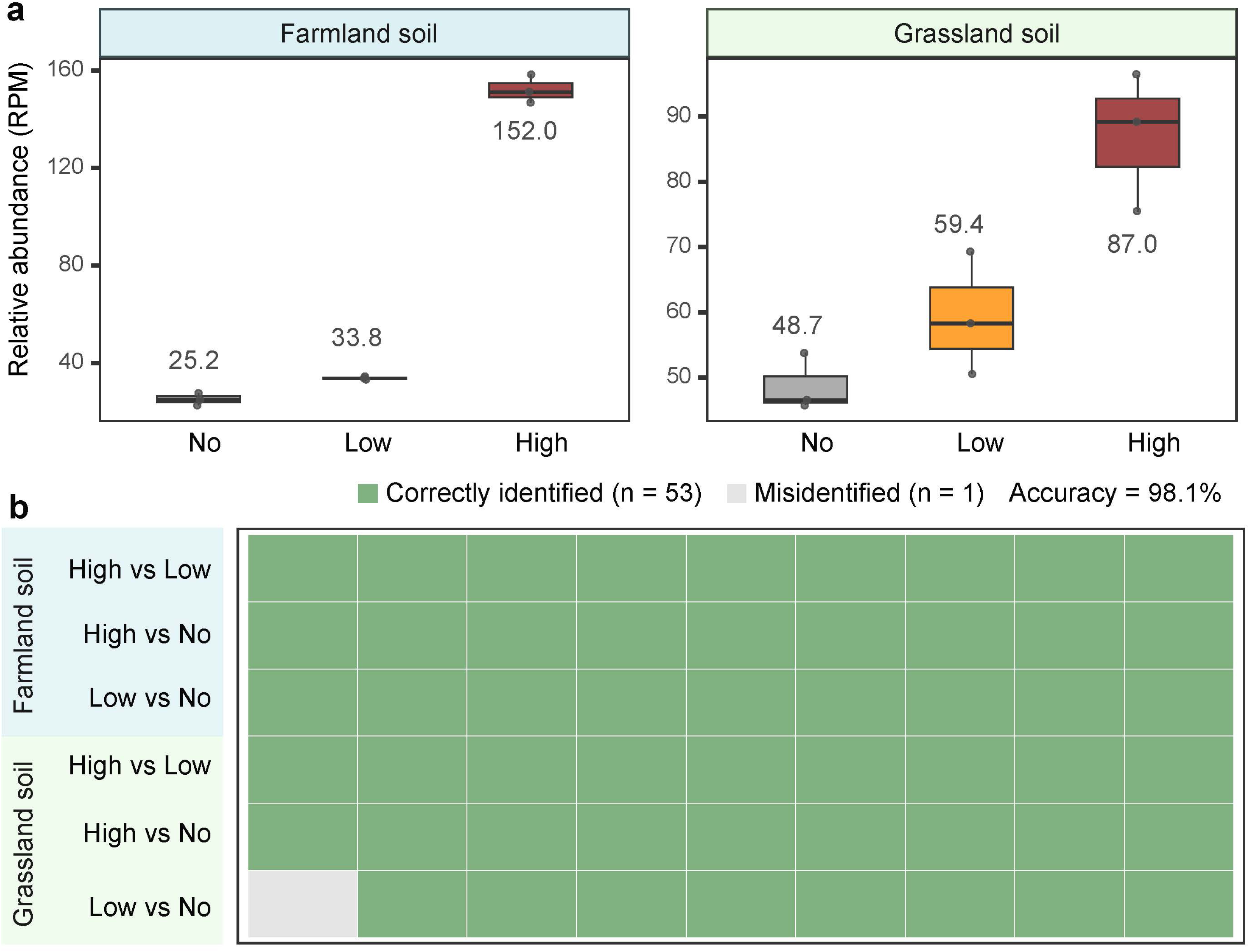
Validation of the accuracy of our pipeline in identifying bacterial pathogens through pathogen DNA spike-in experiments. **a.** Box plots showing that our pipeline can accurately identify pathogens at various concentrations in two different soil types in pathogen DNA spike-in experiments (n = 18). No, no pathogen addition (i.e., control); Low, low pathogen concentration addition; High, high pathogen concentration addition. In box plots, the edges of the box show lower and upper quartiles, the middle line indicates the median. **b.** Using a simplified approach, we evaluate the accuracy of our pipeline by performing pairwise comparisons across all samples treated at different concentrations (with three replicates per concentration). For instance, “High vs Low” involves comparing each of the three samples treated under “High pathogen concentration” conditions against each of the three samples treated under “Low pathogen concentration” conditions. If the values in the “High” treated samples exceed those in the “Low”, our pipeline is considered to have accurately identified the pathogen (i.e., Correctly identified, green). Conversely, if this is not the case, it indicates that our pipeline has not accurately identified the pathogen (i.e., Misidentified, grey). Accuracy = Correctly identified / (Correctly identified + Misidentified).

**Table 1.**
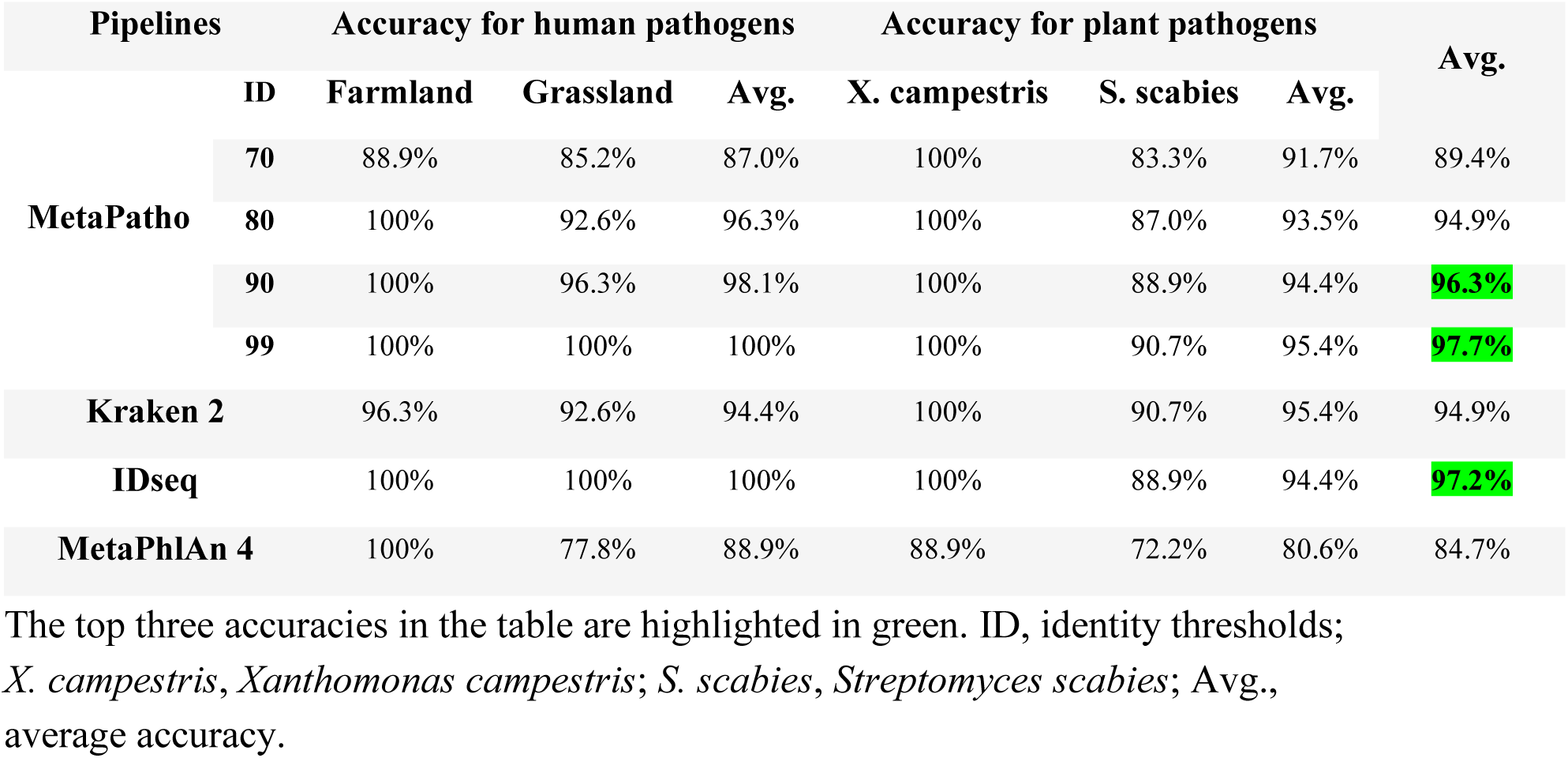
Performance of our tool and other software in pathogen identification.

### Performance validation using pathogen cell spike-in experiments

We further employed pathogen cell spike-in experiments to examine the robustness of our pipeline. We directly introduced cultures of two important plant pathogens (*Xanthomonas campestris* and *Streptomyces scabies*) into soil to prepare soil samples with gradients of pathogen abundance (Fig. 2). Metagenomic analysis results indicated that our pipeline could precisely detect plant pathogens of varying abundances in soil metagenomes (Fig. 5, Table 1). Results from the accuracy calculations suggested that our pipeline achieves an average accuracy of 94.4% (using ID 90) in identifying these two plant pathogens in soil metagenomes. This includes a 100% accuracy for identifying pathogen *Xanthomonas campestris* and an 88.9% accuracy for identifying pathogen *Streptomyces scabies* (Fig. 5, Table 1). For pathogen *Xanthomonas campestris*, the average relative abundances (RPM) identified were 3.4 (no pathogen addition), 15.4 (low concentration addition), 122.4 (medium concentration addition), and 1400.1 (high concentration addition) (Fig. 5). For pathogen *Streptomyces scabies*, the average relative abundances (RPM) identified were 89.7 (no pathogen addition), 91.8 (low concentration addition), 183.6 (medium concentration addition), and 1272.5 (high concentration addition) (Fig. 5). Our analyses showed that at a higher ID threshold (99%), the pipeline achieved an accuracy of up to 95.4% (Extended Data Fig. 2, Table 1). When using a moderate ID threshold (80%), the pipeline maintained a high accuracy of 93.5% (Extended Data Fig. 2, Table 1), supporting accuracy in pathogen indemnification by our tool. Similar to the pathogen DNA spike-in experiments, using lower ID thresholds (60% and 70%) reduced the accuracy of our pipeline to 90-91% (Table 1). Likewise, when identifying plant bacterial pathogens in environmental samples, selecting an appropriate ID threshold (i.e., 80-90%) can improve detection sensitivity while maintaining high accuracy.

**Fig. 5.**
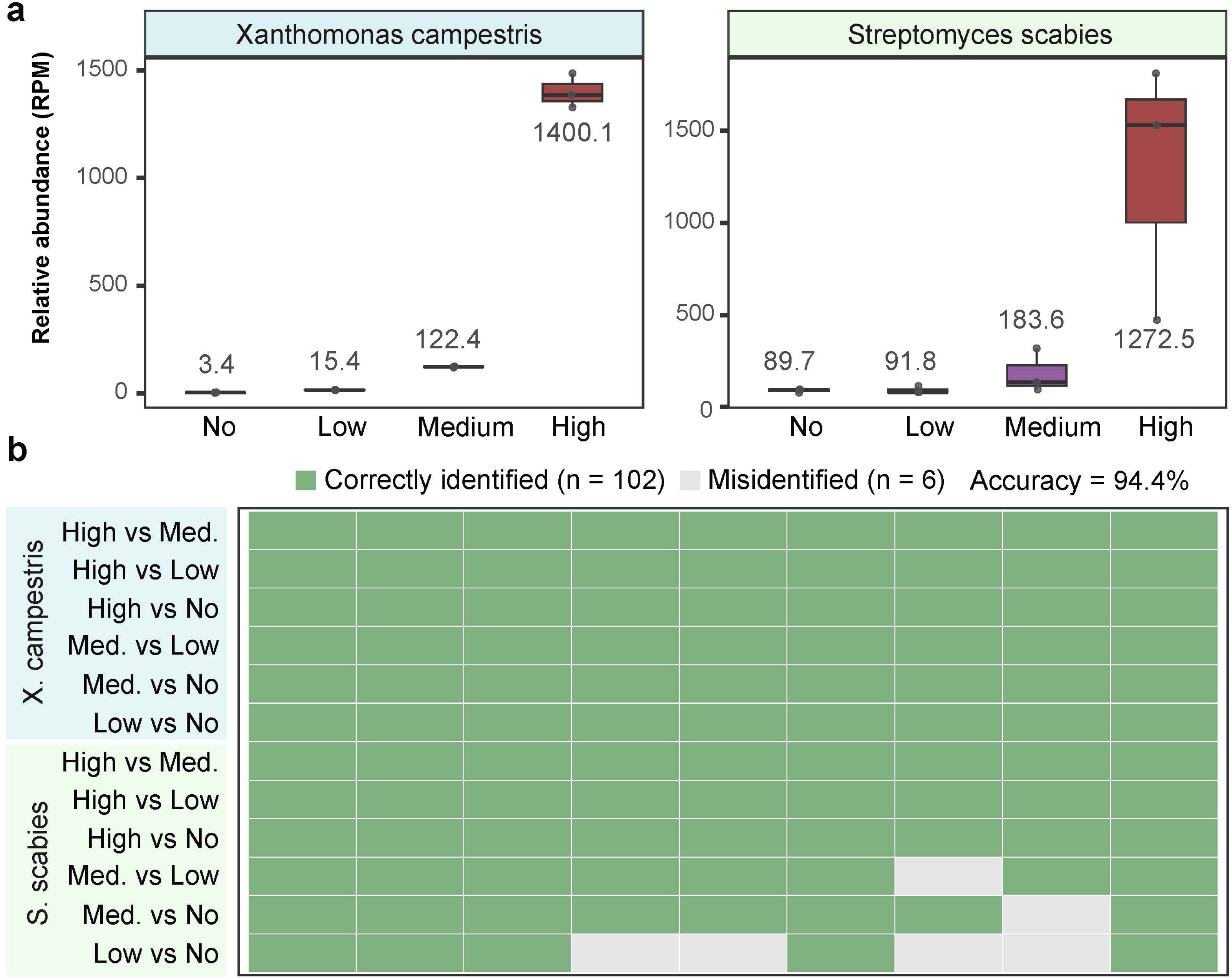
Validation of the accuracy of our pipeline in identifying bacterial pathogens through cell spike-in tests experiments. **a.** Box plots showing that our pipeline can accurately identify various concentrations of two pathogens in pathogen cell spike-in experiments (n = 12). No, no pathogen addition (i.e., control); Low, low pathogen concentration addition; Medium, med., medium pathogen concentration addition; High, high pathogen concentration addition. In box plots, the edges of the box show lower and upper quartiles, the middle line indicates the median. **b.** Using a simplified approach, we evaluate the accuracy of our pipeline by performing pairwise comparisons across all samples treated at different concentrations (with three replicates per concentration). Accuracy = Correctly identified / (Correctly identified + Misidentified).

### Comparison of the performance with other software and pipelines

We employed Kraken 2, MetaPhlAn 4, and IDseq for taxonomic annotation of our soil metagenomes and compared them with our results, thereby further evaluating the performance of our pipeline in the identification of environmental pathogens (Table 1). In pathogen DNA spike-in experiments, Kraken 2 demonstrated an accuracy of 96.3% for pathogen surveillance in farmland soil and 92.6% in grassland soil, with an average accuracy of 94.4% (Extended Data Fig. 3, Table 1). IDseq showed an accuracy of 100% for pathogen surveillance in farmland soil and 100% in grassland soil, with an average accuracy of 100% (Extended Data Fig. 4, Table 1). Moreover, MetaPhlAn 4 demonstrated an accuracy of 100% for pathogen surveillance in farmland soil and 77.8% in grassland soil, with an average accuracy of 88.9% (Extended Data Fig. 5, Table 1). In pathogen cell spike-in experiments, Kraken 2 achieved an accuracy of 100% for *Xanthomonas campestris* and 90.7% for *Streptomyces scabies*, resulting in an average accuracy of 95.4% (Extended Data Fig. 3, Table 1). IDseq showed an accuracy of 100% for *Xanthomonas campestris* and 88.9% for *Streptomyces scabiei*, with an average accuracy of 94.4% (Extended Data Fig. 4, Table 1). In addition, MetaPhlAn 4 demonstrated an accuracy of 88.9% for *Xanthomonas campestris* and 72.2% for *Streptomyces scabies*, with an average accuracy of 80.6% (Extended Data Fig. 5, Table 1).

For the two methods of validation, Kraken 2 and IDseq achieved an average accuracy of 94.9% and 97.2% in pathogen identification, respectively (Table 1). MetaPhlAn 4 achieved an average accuracy of 84.7% in pathogen identification (Table 1). Meanwhile, when applying more stringent ID thresholds (90% and 99%), our pipeline MetaPatho achieved higher pathogen identification accuracies of 96.3% and 97.7% (Table 1), respectively. These findings demonstrate that our pipeline exhibits superior pathogen identification performance. Importantly, compared with other available tools, our pathogen identification pipeline is fully integrated into standard metagenomic analysis workflows and does not require additional software installation or specific environment dependencies. This approach maintains high accuracy while being simple, fast, and user-friendly. For example, when analyzing large sample batches, our pipeline only requires an additional annotation step against the pathogen database on top of standard metagenomic analyses, enabling simultaneous acquisition of pathogen distribution and microbiome functional profiles, such as genes involved in pathogen promotion and suppressions or involved with soil nutrient cycling^50–54^. Such comprehensive information can be harnessed to predict disease and farm productivity outcomes and can support the assessment of soil health and ecosystem restoration.

### Conclusions

Overall, we present a metagenomics-based pipeline, MetaPatho, for identifying human and plant bacterial pathogens in soil metagenomes. In-silico tests based on MOCK community suggested that our pipeline could accurately recognize pathogenic sequences, achieving an overall accuracy of 97.4%. Further results from pathogen DNA and cell spike-in experiments showed that our pipeline can accurately detect pathogen DNA at different concentrations from two soil types, achieving an average accuracy of 96.3%. These combined approaches demonstrate the high accuracy and robustness of our tool in identifying bacterial pathogens in environmental metagenomes. Our pipeline offers crucial tools and data that provides early intelligence on potential public health and food security threats and could supports improved surveillance, health risk management and improved outcomes from the One Health approach.

## Methods

### In-silico tests

To validate our database and pipeline for accurately identifying sequences of bacterial pathogens, we conducted in-silico tests to demonstrate the robustness of our methods. We first constructed a simulated dataset, termed a MOCK community, by randomly selecting translated protein sequences of various lengths (200, 400, 600, and 1000 bp) from the genomes of 40 bacterial pathogens (acting as true positives) and 10 non-pathogenic bacteria (acting as true negatives) (Fig. 2). This resultant MOCK community consisted of approximately 140,000 sequences, including about 100,000 sequences of 200 bp, 25,000 sequences of 400 bp, 10,000 sequences of 600 bp, and 500 sequences of 1000 bp, reflecting the typical length distribution of open reading frames in metagenomic data. We then applied our pipeline to detect human and plant pathogens within the MOCK community.

### Pathogen spike-in experiments

We employed soil spike-in experiments, encompassing both pathogen DNA spike-in and cell spike-in tests, to provide evidence supporting the robustness of our pipeline used in environmental samples. For the pathogen DNA spike-in experiments, we prepared soil DNA samples containing human bacterial pathogens at varying abundances. We initially extracted microbial DNA of two types of soil (farmland and grassland soils) and then added varying concentrations of pathogen genomic DNA (*Mycobacterium tuberculosis* H37Ra, ATCC 25177) to these DNA samples. This method established a gradient of pathogen DNA-containing samples, including DNA samples with no added pathogen, with low concentration of pathogen (2 x 10^4^ copies), and with high (2 x 10^5^ copies) concentration of pathogen (n = 18, three treatments×two soil types×three replicates).

For pathogen cell spike-in experiments, we prepared soil samples containing plant bacterial pathogens at varying abundances. In this experiment, we directly introduced cultures of two important plant pathogens (*Xanthomonas campestris* and *Streptomyces scabies*) into grassland soils to prepare soil samples with gradients of pathogen abundance. This resulted in soil samples with a range of pathogen levels, including no addition, low (1 x 10^4^ CFU), medium (2 x 10^5^ CFU), and high (2 x 10^6^ CFU) concentrations of pathogens (n = 12, four treatments × three replicates). Subsequent to this, DNA was extracted from these soil samples and subjected to shotgun metagenomic sequencing.

### DNA extraction and shotgun sequencing

In soil spike-in experiments, microbial DNA was extracted from 0.3 g of soil samples utilizing the DNeasy PowerSoil Pro Kits (QIAGEN, USA), following the manufacturer’s protocol. The extracted DNA’s quality was assessed via a NanoDrop 2000 spectrophotometer (Thermo Scientific, USA). Shotgun metagenomic sequencing was performed at the Next Generation Sequencing (NGS) Facility of Western Sydney University, Australia. This sequencing utilized the Illumina NovaSeq platform, generating approximately 10 Gb of 150 bp paired-end high-quality reads per sample.

### Bioinformatic analysis

We developed a bioinformatic pipeline to identify human and plant bacterial pathogens from environmental and clinical metagenomes (Fig. 1). Raw sequences were quality-filtered using fastp (v0.20.1)^55^, followed by assembly with Megahit (v1.2.9)^56^, retaining only contigs of ≥1000 bp. High-quality reads were then assembled using Megahit (v1.2.9). The open reading frames (ORFs, i.e., protein-coding genes) of the contigs were predicted using Prodigal (v2.6.3)^57^. These ORFs were clustered at 95% similarity using MMseqs2 (v15.6f452)^58^ to generate a non-redundant gene catalog. Next, taxonomic information of soil-inhabiting bacterial pathogens was annotated by aligning the translated protein sequences from our soil metagenomic datasets against the customized human (HBPDB) and plant (PBPDB) bacterial pathogens database using DIAMOND^44^ (e-value threshold of 1 × 10^−10^, identity > 80%, coverage > 70%, and a minimum alignment length of 100 amino acids). The relative abundance (reads per million, RPM) of each pathogenic phylotype across all samples was determined using Salmon (v1.10.3)^59^. Bacterial virulence factors were annotated by aligning translated protein sequences against the Virulence Factor Database (VFDB)^37^ and the Pathogen–Host Interactions database (PHI-base)^38^ using DIAMOND^44^ (e-value threshold of 1 × 10^−10^, identity > 80%, coverage > 70%). Taxonomic information of human and plant bacterial pathogens from environmental metagenomes at the reads level was also profiled using a combination of Kraken 2 (v2.1.1)^31^ and Bracken (v2.6.2)^60^ tools based on Kraken database (standard Kraken database as of October 2023). In addition, we used MetaPhlAn 4^61^ and IDseq^35^ to perform read-level taxonomic annotation of our soil metagenomes, allowing us to compare the performance of different tools in pathogen identification.

### Calculation of accuracy, specificity, and sensitivity

In the in-silico tests, various performance metrics of our pipeline, including accuracy, specificity, and sensitivity, were calculated following methods described in previously published studies^62,63^. Among these metrics, accuracy (ACC) measures the overall correctness of the pipeline; specificity (SP) measures the proportion of actual negatives that are correctly identified; sensitivity (SE) measures the proportion of actual positives that are correctly identified; positive predictive value (PPV) measures the proportion of positive identifications that are actually correct; and negative predictive value (NPV) measures the proportion of negative identifications that are actually correct. These metrics were calculated using the following formulas: ACC = (TP+TN) / (TP+TN+FP+FN); SP = TN / (TN+FP); SE = TP / (TP+FN); PPV = TP / (TP+FP); NPV = TN / (TN+FN). TP, true positive number; TN, true negative number; FP, false positive number; FN, false negative number.

In soil spike-in experiments, we used a simplified and customized approach to evaluate the accuracy of our pipeline by performing pairwise comparisons across all samples treated at different concentrations (with three replicates per concentration). For instance, “High vs Low” involves comparing each of the three samples treated under “High pathogen concentration” conditions against each of the three samples treated under “Low pathogen concentration” conditions. If the values in the “High” treated samples exceed those in the “Low”, our pipeline is considered to have accurately identified the pathogen (i.e., Correctly identified, green). Conversely, if this is not the case, it indicates that our pipeline has not accurately identified the pathogen (i.e., Misidentified, grey). Finally, the accuracy was calculated using the following formulas: Accuracy = Correctly identified / (Correctly identified + Misidentified).

### Statistical anlyses

The taxonomic table identified from Kraken 2 was rarefied to 700,000 reads per sample for abundance estimates. In the in-silico tests, differences in performance metrics (e.g., accuracy, specificity, and sensitivity) across different ID thresholds were assessed for significance using nonparametric statistical tests (Kruskal-Wallis test). All statistical analyses were carried out in R (http://www.r-project.org).

## Data availability

All raw sequencing data generated in this study have been submitted to the NCBI Sequence Read Archive (SRA) database under the accession numbers PRJNA1290713. The database developed in this study, including the customized human (MetaPatho-HBPDB) and plant (MetaPatho-PBPDB) bacterial pathogen databases, have been deposited at the Figshare (https://figshare.com/s/753d3906f6faf1986d6a) and the Github (https://github.com/ChaoXiong-2024/MetaPatho).

## Code availability

The code used for MetaPatho pipeline and the analyses performed in this study is accessible at the Github (https://github.com/ChaoXiong-2024/MetaPatho) and Figshare (https://figshare.com/s/753d3906f6faf1986d6a).

## Acknowledgments

This work is supported by Australian Research Council Discovery grant (DP2301101448), CRC Future Food Systems, the Department of Agriculture, Fisheries and Forestry (DAFF) Soil Science Challenge grants (ID: 4-H4SSYXD; 4-H4T24R2) to BKS.

## Author contributions

C.X. and B.K.S. conceived and designed the study and the metagenomic pipeline. Laboratory analyses were done by C.X. and M.G. Bioinformatic analyses and statistical analyses were done by C.X. and B.K.S. The manuscript was written by C.X. and B.K.S., with contributions from M.G.

## Additional information

Supplementary Information is available for this paper.

**Correspondence and requests for materials** should be addressed to Brajesh K. Singh.

## Extended data figures and tables

**Extended Data Fig. 1 | Validation of the accuracy of our pipeline (using ID 99 and ID 80) in identifying bacterial pathogens through pathogen DNA spike-in experiments. a.** Box plots showing that our pipeline can accurately identify human bacterial pathogens at various concentrations in two different soil types in pathogen DNA spike-in experiments (ID 99, n = 18). **b.** Using a simplified approach, we evaluate the accuracy of our pipeline (ID 99, n = 18) by performing pairwise comparisons across all samples treated at different concentrations (with three replicates per concentration). For instance, “High vs Low” involves comparing each of the three samples treated under “High pathogen concentration” conditions against each of the three samples treated under “Low pathogen concentration” conditions. If the values in the “High” treated samples exceed those in the “Low”, our pipeline is considered to have accurately identified the pathogen (i.e., Correctly identified, green). Conversely, if this is not the case, it indicates that our pipeline has not accurately identified the pathogen (i.e., Misidentified, grey). Accuracy = Correctly identified / (Correctly identified + Misidentified). **c.** Box plots showing that our pipeline can accurately identify human bacterial pathogens at various concentrations in two different soil types in pathogen DNA spike-in experiments (ID 80, n = 18). **b.** Using a simplified approach, we evaluate the accuracy of our pipeline (ID 80, n = 18) by performing pairwise comparisons across all samples treated at different concentrations (with three replicates per concentration). Accuracy = Correctly identified / (Correctly identified + Misidentified). No, no pathogen addition (i.e., control); Low, low pathogen concentration addition; High, high pathogen concentration addition. In box plots, the edges of the box show lower and upper quartiles, the middle line indicates the median.

**Extended Data Fig. 2 | Validation of the accuracy of our pipeline (using ID 99 and ID 80) in identifying bacterial pathogens through cell spike-in tests experiments. a.** Box plots showing that our pipeline can accurately identify various concentrations of plant bacterial pathogens in pathogen cell spike-in experiments (ID 99, n = 12). **b.** Using a simplified approach, we evaluate the accuracy of our pipeline (ID 99, n = 12) by performing pairwise comparisons across all samples treated at different concentrations (with three replicates per concentration). Accuracy = Correctly identified / (Correctly identified + Misidentified). **c.** Box plots showing that our pipeline can accurately identify various concentrations of plant bacterial pathogens in pathogen cell spike-in experiments (ID 80, n = 12). **d.** Using a simplified approach, we evaluate the accuracy of our pipeline (ID 80, n = 12) by performing pairwise comparisons across all samples treated at different concentrations (with three replicates per concentration). Accuracy = Correctly identified / (Correctly identified + Misidentified). No, no pathogen addition (i.e., control); Low, low pathogen concentration addition; Medium, med., medium pathogen concentration addition; High, high pathogen concentration addition. In box plots, the edges of the box show lower and upper quartiles, the middle line indicates the median.

**Extended Data Fig. 3 | The performance of Kraken 2 in identifying bacterial pathogens in pathogen DNA spike-in experiments and cell spike-in tests experiments. a.** Box plots showing the performance of the Kraken 2 in identifying human bacterial pathogens at various concentrations in two different soil types in pathogen DNA spike-in experiments (n = 18). **b.** Using a simplified approach, we evaluate the accuracy of the Kraken 2 in identifying human bacterial pathogens by performing pairwise comparisons across all samples treated at different concentrations (with three replicates per concentration). Accuracy = Correctly identified / (Correctly identified + Misidentified). **c.** Box plots showing the performance of the Kraken 2 in identifying plant bacterial pathogens in pathogen cell spike-in experiments (n = 12). **b.** Using a simplified approach, we evaluate the accuracy of the Kraken 2 in identifying plant bacterial pathogens by performing pairwise comparisons across all samples treated at different concentrations (with three replicates per concentration). Accuracy = Correctly identified / (Correctly identified + Misidentified). No, no pathogen addition (i.e., control); Low, low pathogen concentration addition; Medium, med., medium pathogen concentration addition; High, high pathogen concentration addition. In box plots, the edges of the box show lower and upper quartiles, the middle line indicates the median.

**Extended Data Fig. 4 | The performance of IDseq in identifying bacterial pathogens in pathogen DNA spike-in experiments and cell spike-in tests experiments. a.** Box plots showing the performance of the IDseq in identifying human bacterial pathogens at various concentrations in two different soil types in pathogen DNA spike-in experiments (n = 18). **b.** Using a simplified approach, we evaluate the accuracy of the IDseq in identifying human bacterial pathogens by performing pairwise comparisons across all samples treated at different concentrations (with three replicates per concentration). Accuracy = Correctly identified / (Correctly identified + Misidentified). **c.** Box plots showing the performance of the IDseq in identifying plant bacterial pathogens in pathogen cell spike-in experiments (n = 12). **b.** Using a simplified approach, we evaluate the accuracy of the IDseq in identifying plant bacterial pathogens by performing pairwise comparisons across all samples treated at different concentrations (with three replicates per concentration). Accuracy = Correctly identified / (Correctly identified + Misidentified). No, no pathogen addition (i.e., control); Low, low pathogen concentration addition; Medium, med., medium pathogen concentration addition; High, high pathogen concentration addition. In box plots, the edges of the box show lower and upper quartiles, the middle line indicates the median.

**Extended Data Fig. 5 | The performance of MetaPhlAn 4 in identifying bacterial pathogens in pathogen DNA spike-in experiments and cell spike-in tests experiments. a.** Box plots showing the performance of the MetaPhlAn 4 in identifying human bacterial pathogens at various concentrations in two different soil types in pathogen DNA spike-in experiments (n = 18). **b.** Using a simplified approach, we evaluate the accuracy of the MetaPhlAn 4 in identifying human bacterial pathogens by performing pairwise comparisons across all samples treated at different concentrations (with three replicates per concentration). Accuracy = Correctly identified / (Correctly identified + Misidentified). **c.** Box plots showing the performance of the MetaPhlAn 4 in identifying plant bacterial pathogens in pathogen cell spike-in experiments (n = 12). **d.** Using a simplified approach, we evaluate the accuracy of the MetaPhlAn 4 in identifying plant bacterial pathogens by performing pairwise comparisons across all samples treated at different concentrations (with three replicates per concentration). Accuracy = Correctly identified / (Correctly identified + Misidentified). No, no pathogen addition (i.e., control); Low, low pathogen concentration addition; Medium, med., medium pathogen concentration addition; High, high pathogen concentration addition. In box plots, the edges of the box show lower and upper quartiles, the middle line indicates the median.

**Extended Data Table 1** List of human bacterial pathogens in the MetaPatho-HBPDB

**Extended Data Table 2** List of plant bacterial pathogens in the MetaPatho-PBPDB

