## Extended Data Figures and Tables for "A metagenomics-based tool for surveillance of bacterial pathogens in environmental samples"

Ph: +61 2 4570 1329 | Mob: +61 404187962

### Extended Data Figures

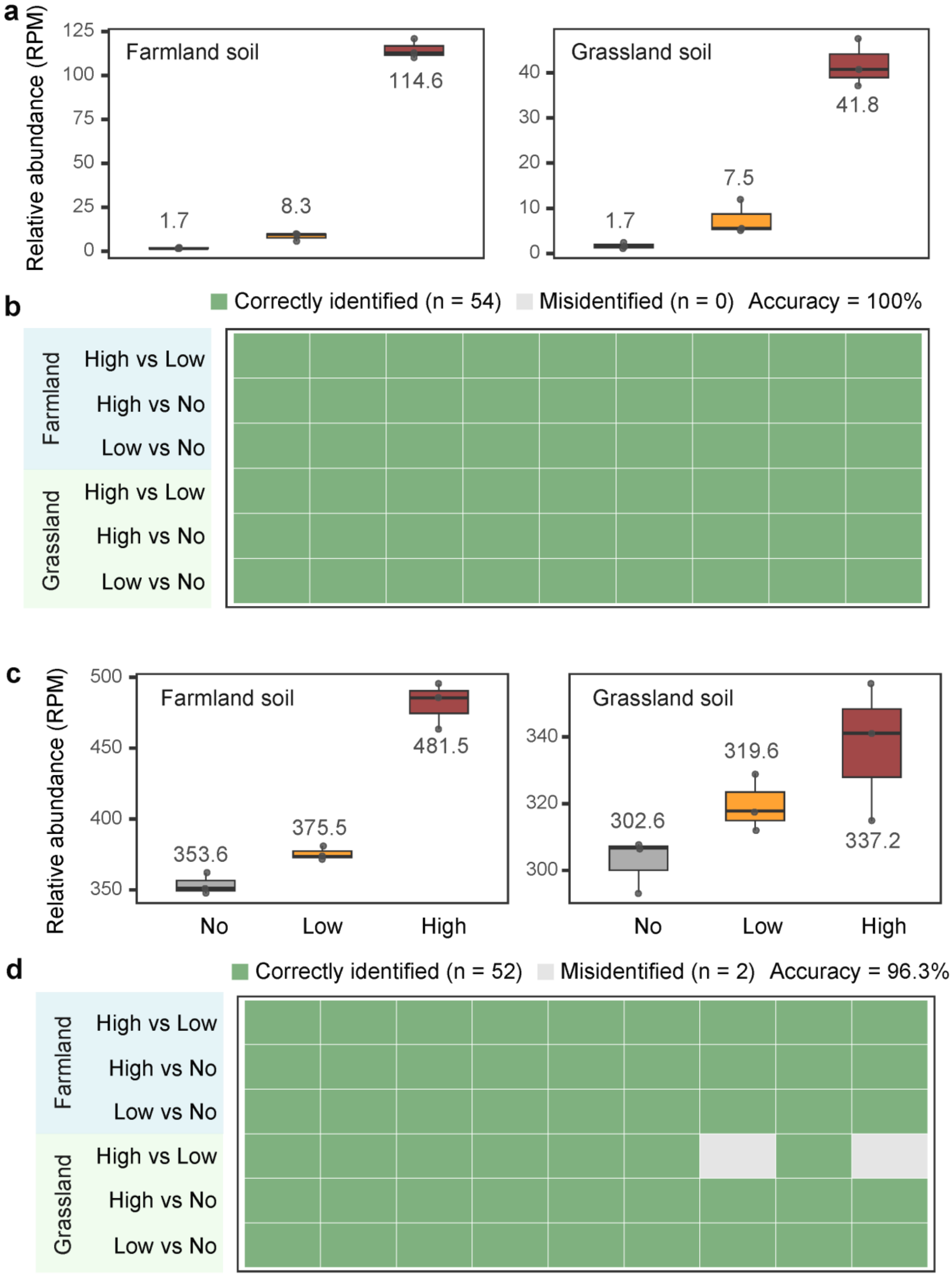

**Extended Data Fig. 1 | Validation of the accuracy of our pipeline (using ID 99 and ID 80) in identifying bacterial pathogens through pathogen DNA spike-in experiments. a.** Box plots showing that our pipeline can accurately identify human bacterial pathogens at various concentrations in two different soil types in pathogen DNA spike-in experiments (ID 99, n = 18). **b.** Using a simplified approach, we evaluate the accuracy of our pipeline (ID 99, n = 18) by performing pairwise comparisons across all samples treated at different concentrations (with three replicates per concentration). For instance, "High vs Low" involves comparing each of the three samples treated under "High pathogen concentration" conditions against each of the three samples treated under "Low pathogen concentration" conditions. If the values in the "High" treated samples exceed those in the "Low", our pipeline is considered to have accurately identified the pathogen (i.e., Correctly identified, green). Conversely, if this is not the case, it indicates that our pipeline has not accurately identified the pathogen (i.e., Misidentified, grey). Accuracy = Correctly identified / (Correctly identified + Misidentified). **c.** Box plots showing that our pipeline can accurately identify human bacterial pathogens at various concentrations in two different soil types in pathogen DNA spike-in experiments (ID 80, n = 18). **b.** Using a simplified approach, we evaluate the accuracy of our pipeline (ID 80, n = 18) by performing pairwise comparisons across all samples treated at different concentrations (with three replicates per concentration). Accuracy = Correctly identified / (Correctly identified + Misidentified). No, no pathogen addition (i.e., control); Low, low pathogen concentration addition; High, high pathogen concentration addition. In box plots, the edges of the box show lower and upper quartiles, the middle line indicates the median.

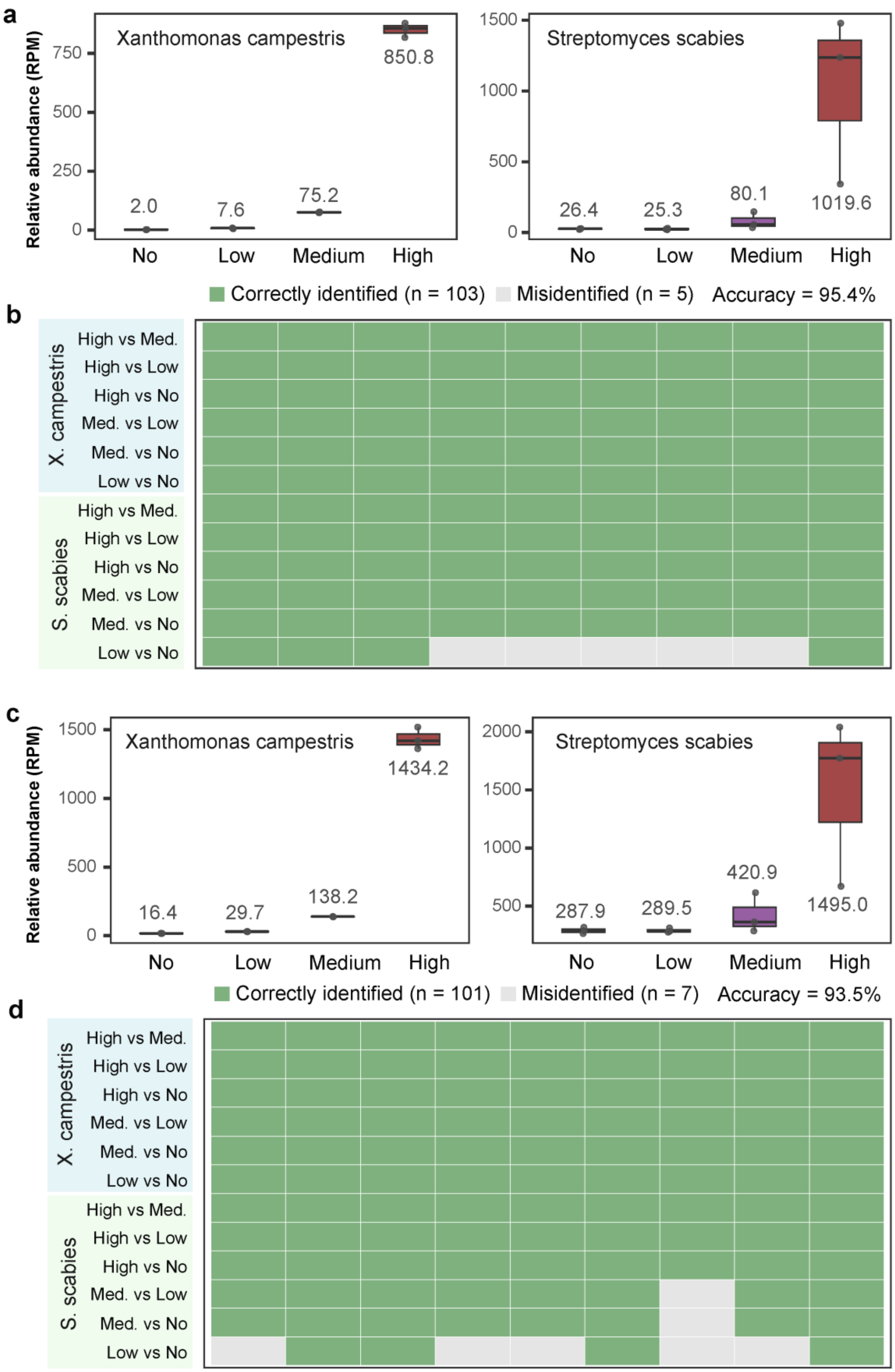

a
Relative abundance (RPM)
Xanthomonas campestris
850.8
2.0
7.6
75.2
Streptomyces scabies
26.4
25.3
80.1
1019.6
No
Low
Medium
High
b
Correctly identified (n = 103)
Misidentified (n = 5)
Accuracy = 95.4%
X. campestris
S. scabies
High vs Med.
High vs Low
High vs No
Med. vs Low
Med. vs No
Low vs No
c
Xanthomonas campestris
1434.2
16.4
29.7
138.2
Streptomyces scabies
287.9
289.5
420.9
1495.0
d
Correctly identified (n = 101)
Misidentified (n = 7)
Accuracy = 93.5%

**Extended Data Fig. 2 | Validation of the accuracy of our pipeline (using ID 99 and ID 80) in identifying bacterial pathogens through cell spike-in tests experiments. a.** Box plots showing that our pipeline can accurately identify various concentrations of plant bacterial pathogens in pathogen cell spike-in experiments (ID 99, n = 12). **b.** Using a simplified approach, we evaluate the accuracy of our pipeline (ID 99, n = 12) by performing pairwise comparisons across all samples treated at different concentrations (with three replicates per concentration). Accuracy = Correctly identified / (Correctly identified + Misidentified). **c.** Box plots showing that our pipeline can accurately identify various concentrations of plant bacterial pathogens in pathogen cell spike-in experiments (ID 80, n = 12). **d.** Using a simplified approach, we evaluate the accuracy of our pipeline (ID 80, n = 12) by performing pairwise comparisons across all samples treated at different concentrations (with three replicates per concentration). Accuracy = Correctly identified / (Correctly identified + Misidentified). No, no pathogen addition (i.e., control); Low, low pathogen concentration addition; Medium, med., medium pathogen concentration addition; High, high pathogen concentration addition. In box plots, the edges of the box show lower and upper quartiles, the middle line indicates the median.

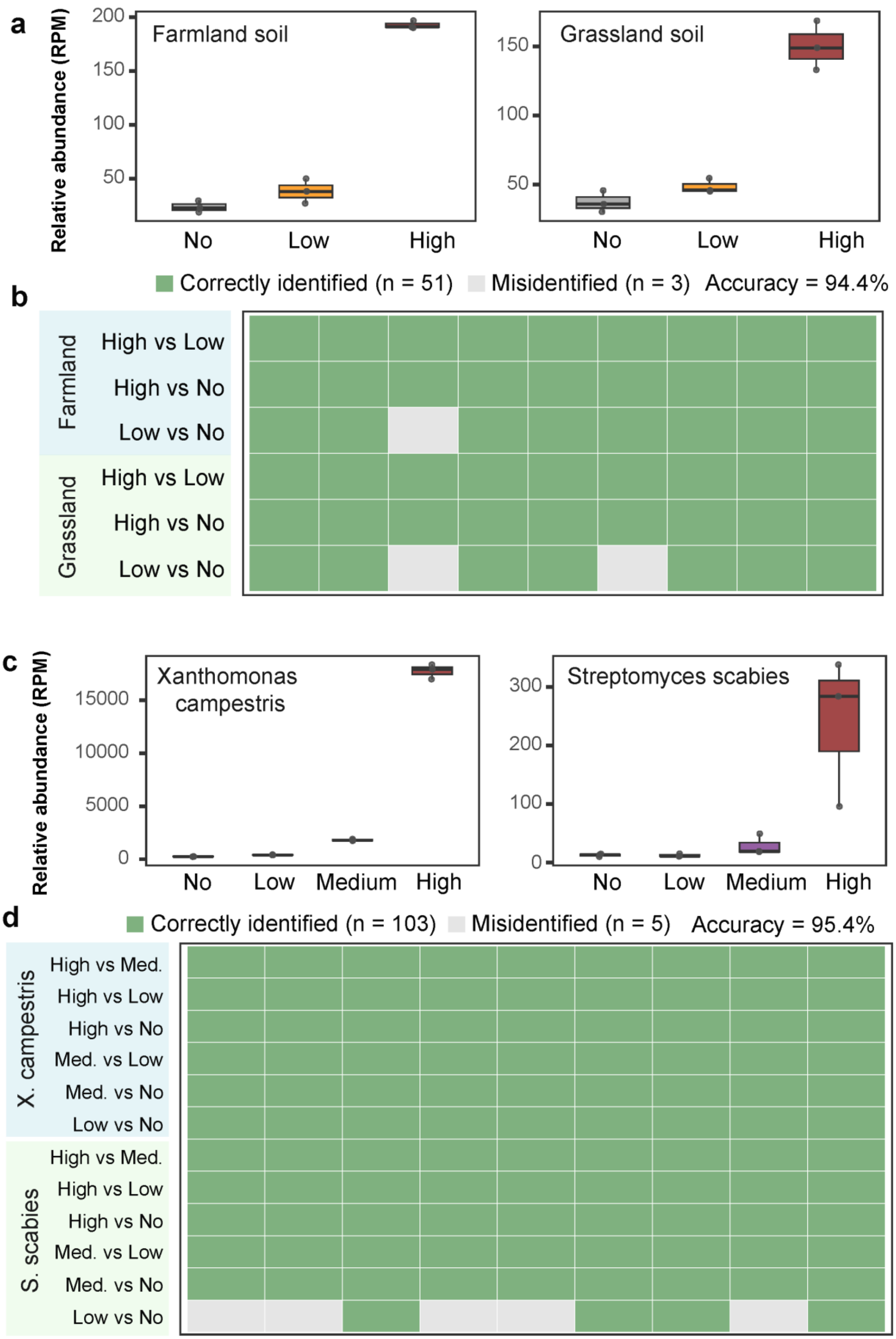
a
Relative abundance (RPM)
Farmland soil
200
150
100
50
No
Low
High
Grassland soil
150
100
50
No
Low
High
b
Correctly identified (n = 51)
Misidentified (n = 3)
Accuracy = 94.4%
Farmland
High vs Low
High vs No
Low vs No
Grassland
High vs Low
High vs No
Low vs No
c
Relative abundance (RPM)
Xanthomonas campestris
15000
10000
5000
0
No
Low
Medium
High
Streptomyces scabies
300
200
100
0
No
Low
Medium
High
d
Correctly identified (n = 103)
Misidentified (n = 5)
Accuracy = 95.4%
X. campestris
High vs Med.
High vs Low
High vs No
Med. vs Low
Med. vs No
Low vs No
S. scabies
High vs Med.
High vs Low
High vs No
Med. vs Low
Med. vs No
Low vs No

**Extended Data Fig. 3 | The performance of Kraken 2 in identifying bacterial pathogens in pathogen DNA spike-in experiments and cell spike-in tests experiments. a.** Box plots showing the performance of the Kraken 2 in identifying human bacterial pathogens at various concentrations in two different soil types in pathogen DNA spike-in experiments (n = 18). **b.** Using a simplified approach, we evaluate the accuracy of the Kraken 2 in identifying human bacterial pathogens by performing pairwise comparisons across all samples treated at different concentrations (with three replicates per concentration). Accuracy = Correctly identified / (Correctly identified + Misidentified). **c.** Box plots showing the performance of the Kraken 2 in identifying plant bacterial pathogens in pathogen cell spike-in experiments (n = 12). **b.** Using a simplified approach, we evaluate the accuracy of the Kraken 2 in identifying plant bacterial pathogens by performing pairwise comparisons across all samples treated at different concentrations (with three replicates per concentration). Accuracy = Correctly identified / (Correctly identified + Misidentified). No, no pathogen addition (i.e., control); Low, low pathogen concentration addition; Medium, med., medium pathogen concentration addition; High, high pathogen concentration addition. In box plots, the edges of the box show lower and upper quartiles, the middle line indicates the median.

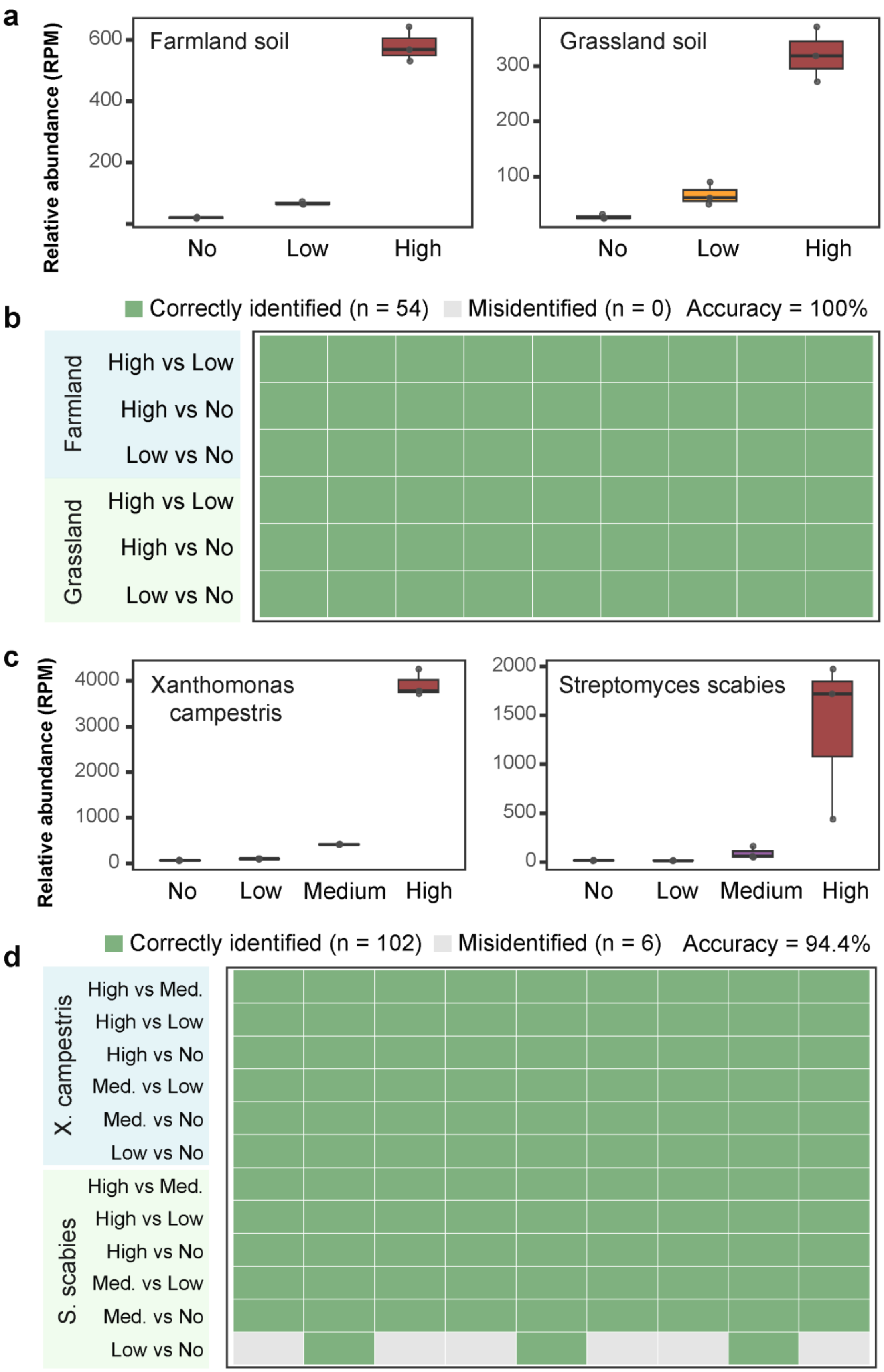
a
Relative abundance (RPM)
Farmland soil
600
400
200
No
Low
High
Grassland soil
300
200
100
No
Low
High
b
Correctly identified (n = 54)
Misidentified (n = 0)
Accuracy = 100%
Farmland
High vs Low
High vs No
Low vs No
Grassland
High vs Low
High vs No
Low vs No
c
Relative abundance (RPM)
Xanthomonas campestris
4000
3000
2000
1000
0
No
Low
Medium
High
Streptomyces scabies
2000
1500
1000
500
0
No
Low
Medium
High
d
Correctly identified (n = 102)
Misidentified (n = 6)
Accuracy = 94.4%
X. campestris
High vs Med.
High vs Low
High vs No
Med. vs Low
Med. vs No
Low vs No
S. scabies
High vs Med.
High vs Low
High vs No
Med. vs Low
Med. vs No
Low vs No

**Extended Data Fig. 4 | The performance of IDseq in identifying bacterial pathogens in pathogen DNA spike-in experiments and cell spike-in tests experiments. a.** Box plots showing the performance of the IDseq in identifying human bacterial pathogens at various concentrations in two different soil types in pathogen DNA spike-in experiments (n = 18). **b.** Using a simplified approach, we evaluate the accuracy of the IDseq in identifying human bacterial pathogens by performing pairwise comparisons across all samples treated at different concentrations (with three replicates per concentration). Accuracy = Correctly identified / (Correctly identified + Misidentified). **c.** Box plots showing the performance of the IDseq in identifying plant bacterial pathogens in pathogen cell spike-in experiments (n = 12). **b.** Using a simplified approach, we evaluate the accuracy of the IDseq in identifying plant bacterial pathogens by performing pairwise comparisons across all samples treated at different concentrations (with three replicates per concentration). Accuracy = Correctly identified / (Correctly identified + Misidentified). No, no pathogen addition (i.e., control); Low, low pathogen concentration addition; Medium, med., medium pathogen concentration addition; High, high pathogen concentration addition. In box plots, the edges of the box show lower and upper quartiles, the middle line indicates the median.

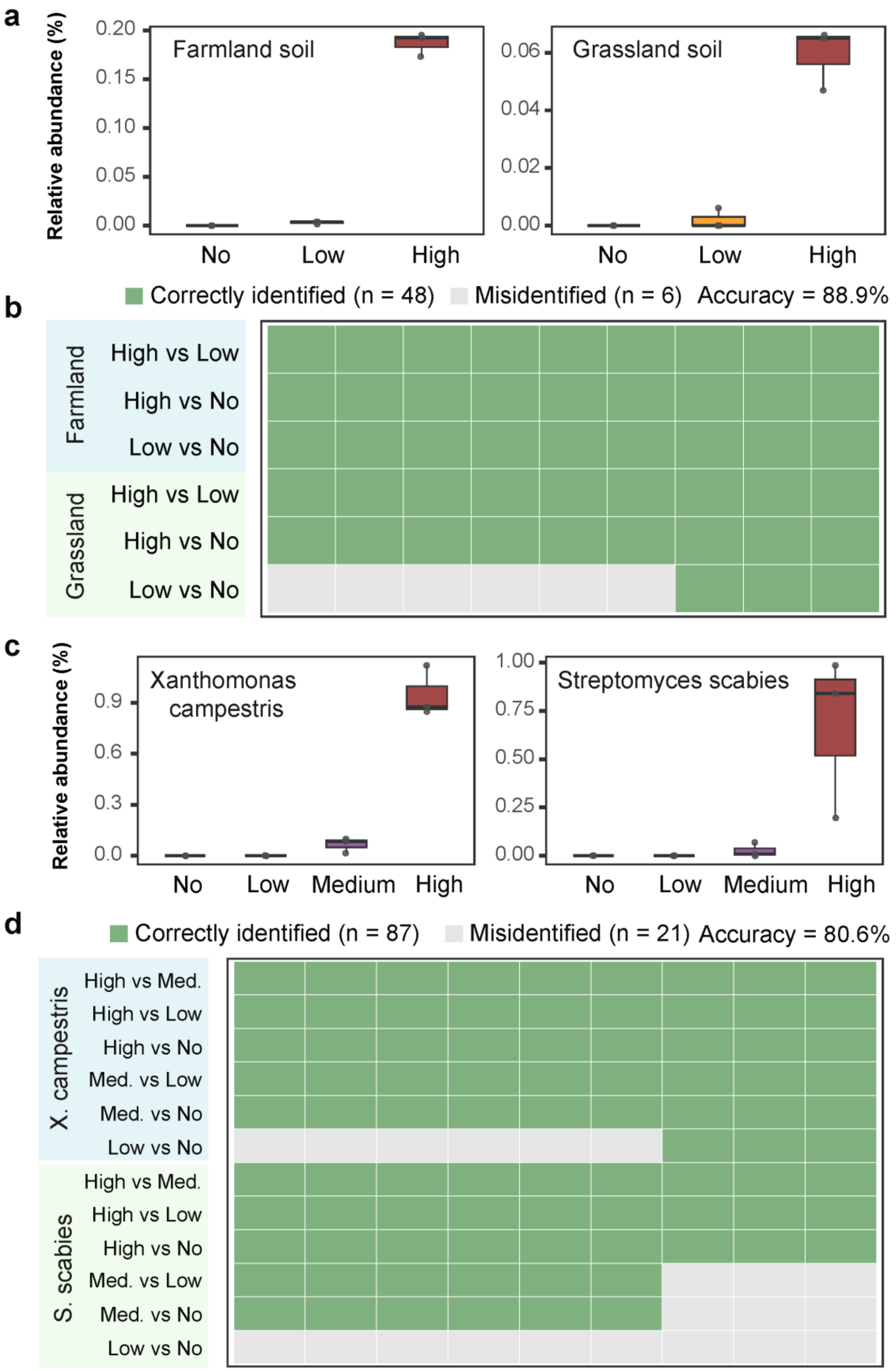
a
Relative abundance (%)
Farmland soil
Grassland soil
0.20
0.15
0.10
0.05
0.00
0.06
0.04
0.02
0.00
No
Low
High
b
Correctly identified (n = 48)
Misidentified (n = 6)
Accuracy = 88.9%
Farmland
Grassland
High vs Low
High vs No
Low vs No
c
Relative abundance (%)
Xanthomonas campestris
Streptomyces scabies
0.9
0.6
0.3
0.0
1.00
0.75
0.50
0.25
0.00
No
Low
Medium
High
d
Correctly identified (n = 87)
Misidentified (n = 21)
Accuracy = 80.6%
X. campestris
S. scabies
High vs Med.
High vs Low
High vs No
Med. vs Low
Med. vs No
Low vs No

**Extended Data Fig. 5 | The performance of MetaPhlAn 4 in identifying bacterial pathogens in pathogen DNA spike-in experiments and cell spike-in tests experiments. a.** Box plots showing the performance of the MetaPhlAn 4 in identifying human bacterial pathogens at various concentrations in two different soil types in pathogen DNA spike-in experiments (n = 18). **b.** Using a simplified approach, we evaluate the accuracy of the MetaPhlAn 4 in identifying human bacterial pathogens by performing pairwise comparisons across all samples treated at different concentrations (with three replicates per concentration). Accuracy = Correctly identified / (Correctly identified + Misidentified). **c.** Box plots showing the performance of the MetaPhlAn 4 in identifying plant bacterial pathogens in pathogen cell spike-in experiments (n = 12). **d.** Using a simplified approach, we evaluate the accuracy of the MetaPhlAn 4 in identifying plant bacterial pathogens by performing pairwise comparisons across all samples treated at different concentrations (with three replicates per concentration). Accuracy = Correctly identified / (Correctly identified + Misidentified). No, no pathogen addition (i.e., control); Low, low pathogen concentration addition; Medium, med., medium pathogen concentration addition; High, high pathogen concentration addition. In box plots, the edges of the box show lower and upper quartiles, the middle line indicates the median.

### Extended Data Tables

#### Extended Data Table 1 List of human bacterial pathogens in the MetaPatho-HBPDB

| Human bacterial pathogens | Human diseases |
|---|---|
| Acinetobacter baumannii | Infections |
| Aeromonas hydrophila | Infections |
| Aeromonas salmonicida | Infections |
| Aeromonas veronii | Gastrointestinal diseases |
| Anaplasma phagocytophilum | Anaplasmosis |
| Bacillus anthracis | Anthrax |
| Bacillus cereus | Food poisoning |
| Bacillus cytotoxicus | Gastrointestinal diseases |
| Bacillus paranthracis | Gastrointestinal diseases |
| Brucella anthropi | Infections |
| Burkholderia ambifaria | Infections |
| Burkholderia cenocepacia | Infections |
| Burkholderia cepacia | Infections |
| Burkholderia pseudomallei | Melioidosis |
| Campylobacter coli | Gastrointestinal diseases |
| Campylobacter fetus | Infections |
| Campylobacter jejuni | Gastrointestinal diseases |
| Campylobacter lari | Gastrointestinal diseases |
| Campylobacter upsaliensis | Gastrointestinal diseases |
| Chlamydia psittaci | Parrot fever |
| Chromobacterium violaceum | Infections |
| Citrobacter braakii | Infections |
| Citrobacter freundii | Infections |
| Citrobacter koseri | Infections |
| Clostridioides difficile | Pseudomembranous colitis and infections |
| Clostridium botulinum | Paralysis |
| Clostridium novyi | Gas gangrene |
| Clostridium perfringens | Food poisoning |
| Clostridium tetani | Tetanus |
| Coxiella burnetii | Q fever |
| Cronobacter sakazakii | Meningitis |
| Edwardsiella tarda | Gastrointestinal diseases |
| Elizabethkingia anophelis | Meningitis |
| Enterobacter cancerogenus | Infections |
| Enterobacter roggenkampii | Gastrointestinal diseases |
| Enterococcus casseliflavus | Infections |
| Enterococcus faecalis | Infections |
| Enterococcus faecium | Infections |
| Escherichia coli | Gastrointestinal diseases |

| | |
|---|---|
| Francisella tularensis | Tularemia |
| Helicobacter pylori | Infections |
| Klebsiella oxytoca | Infections |
| Klebsiella pneumoniae | Respiratory infections |
| Klebsiella variicola | Respiratory infections |
| Kluyvera ascorbata | Infections |
| Legionella longbeachae | Legionnaires' disease |
| Legionella pneumophila | Legionnaires' disease |
| Leptospira interrogans | Leptospirosis |
| Listeria ivanovii | Listeriosis |
| Listeria monocytogenes | Listeriosis |
| Morganella morganii | Infections |
| Mycobacterium leprae | Leprosy |
| Mycobacterium tuberculosis | Tuberculosis |
| Mycobacteroides abscessus | Infections |
| Mycoplasmoides pneumoniae | Atypical pneumonia |
| Neisseria gonorrhoeae | Gonorrhea |
| Neisseria meningitidis | Meningococcal meningitis |
| Nocardia farcinica | Nocardiosis |
| Pantoea agglomerans | Infections |
| Photobacterium damselae | Infections |
| Proteus mirabilis | Infections |
| Providencia alcalifaciens | Gastrointestinal diseases |
| Pseudomonas aeruginosa | Infections |
| Salmonella enterica | Salmonellosis |
| Serratia marcescens | Infections |
| Shigella boydii | Shigellosis |
| Shigella dysenteriae | Shigellosis |
| Shigella flexneri | Shigellosis |
| Shigella sonnei | Shigellosis |
| Staphylococcus aureus | Infections |
| Staphylococcus pseudintermedius | Infections |
| Streptococcus agalactiae | Infections |
| Streptococcus iniae | Infections |
| Streptococcus pneumoniae | Pneumonia, meningitis, and sepsis |
| Streptococcus pyogenes | Strep throat, rheumatic fever, and skin infections |
| Streptococcus suis | Infections |
| Vibrio cholerae | Cholera |
| Vibrio parahaemolyticus | Gastrointestinal diseases |
| Vibrio vulnificus | Wound infections and primary septicemia |
| Yersinia enterocolitica | Yersiniosis |
| Yersinia pestis | Plague |
| Yersinia pseudotuberculosis | Infections |

**Extended Data Table 2 List of plant bacterial pathogens in the MetaPatho-PBPDB**

| Plant bacterial pathogens | Plant Diseases |
|---|---|
| Agrobacterium larrymoorei | Crown Gall Disease |
| Agrobacterium rubi | Crown and Root Gall, Hairy Root |
| Agrobacterium tumefaciens | Crown Gall Disease |
| Agrobacterium vitis | crown gall disease |
| Brenneria goodwinii | oak and oriental beech decline |
| Brenneria nigrifluens | Bacterial Canker, Shallow-Bark Canker |
| Brenneria rubrifaciens | Deep Bark Canker Disease |
| Burkholderia gladioli | Leaf spots and stripe, Bacterial Crown Rot, Panicle Blight |
| Burkholderia glumae | Bacterial Panicle Blight |
| Burkholderia plantarii | Rice Seedling Blight |
| Candidatus Liberibacter africanus | Huanglongbing, citrus greening disease |
| Candidatus Liberibacter americanus | Citrus Huanglongbing |
| Candidatus Liberibacter asiaticus | Citrus Huanglongbing |
| Candidatus Phytoplasma asteris | aster yellows |
| Candidatus Phytoplasma rubi | Rubus Stunt Disease |
| Candidatus Liberibacter solanacearum | Overall chlorosis, severe stunting, leaf cupping |
| Candidatus Phytoplasma mali | Apple proliferation disease |
| Candidatus Phytoplasma solani | Black wood, Rubbery Taproot Disease |
| Candidatus Phytoplasma ziziphi | Yellowing leaves, Broom Disease |
| Clavibacter michiganensis | Canker |
| Clavibacter sepedonicus | Wilt and tuber rot |
| Curtobacterium flaccumfaciens | Silvering disease |
| Dickeya dadantii | soft rot, brown rot or blackleg |
| Dickeya dianthicola | Soft rot, brown rot or blackleg |
| Dickeya fangzhongdai | Soft Rot Disease |
| Dickeya solani | Blackleg |
| Dickeya zeae | Bacterial Foot Rot, Soft Rot Disease |
| Dickeya chrysanthemi | Bacterial Soft Rot |
| Enterobacter mori | Wilt, soft rot, Leaf Spot |
| Erwinia amylovora | Fire blight |
| Erwinia persicina | Stalk Rot, Leaf Spot, Pink Disease, Bulb Rot |
| Erwinia pyrifoliae | Necrotic diseas |
| Erwinia rhapontici | Crown and Shoot Rot, pink seed, Leaf Spot, Stalk Rot |
| Erwinia tracheiphila | Wilt |
| Leifsonia xyli | Ratoon Stunting Disease |
| Lonsdalea populi | Bark Canker Disease |
| Pantoea ananatis | Crown Necrobiosis, Blight, Stem rot, leaf streak, leaf spot |
| Pantoea stewartii | Stewart's wilt and blight |
| Paracidovorax avenae | Bacterial Leaf Stripe |
| Paracidovorax cattleyae | Bacterial Brown Spot |
| Paracidovorax citrulli | Seedling blight and bacterial fruit blotch |

| | |
|---|---|
| Paracidovorax oryzae | Rice bacterial brown streak disease |
| Pectobacterium actinidiae | Canker |
| Pectobacterium aroidearum | soft rot |
| Pectobacterium atrosepticum | Blackleg Disease, Stalk and Head Rot, Soft Rot |
| Pectobacterium brasiliense | Blackleg and Soft Rot |
| Pectobacterium cacticida | Soft Rot |
| Pectobacterium carotovorum | Soft Rot |
| Pectobacterium colocasium | Soft Rot |
| Pectobacterium odoriferum | Soft Rot |
| Pectobacterium parmentieri | Blackleg, Soft Rot, stem rot |
| Pectobacterium parvum | Stem rot |
| Pectobacterium polaris | Soft Rot, Stem rot, Blackleg |
| Pectobacterium punjabense | Blackleg and Soft Rot |
| Pectobacterium versatile | Blackleg, Soft Rot, stem rot |
| Pectobacterium wasabiae | Soft Rot, stem rot |
| Pseudomonas amygdali | Bacterial Gall |
| Pseudomonas avellanae | Canker |
| Pseudomonas cannabina | leaf and stem rot, Blight |
| Pseudomonas cerasi | shoot blight |
| Pseudomonas cichorii | Blight, Leaf Spot |
| Pseudomonas coronafaciens | Leaf and stem rot of hemp, Halo Blight |
| Pseudomonas corrugata | Pith Necrosis |
| Pseudomonas fuscovaginae | Brown sheath rot |
| Pseudomonas marginalis | Soft rot and marginal leaf necrosis |
| Pseudomonas palleroniana | Soft Rot |
| Pseudomonas psychrotolerans | Leaf Spot |
| Pseudomonas savastanoi | Canker, Knot |
| Pseudomonas syringae | Blight, Rot, Leaf spot |
| Pseudomonas tremae | Gall disease |
| Pseudomonas viridiflava | Necrotic lesions, basal stem and root rots, pith necrosis |
| Pseudomonas mediterranea | pith necrosis |
| Ralstonia pseudosolanacearum | Wilt |
| Ralstonia solanacearum | Wilt |
| Ralstonia syzygii | Wilt |
| Rathayibacter rathayi | Gumming disease, Head Blight |
| Rathayibacter tritici | spike blight |
| Rathayibacter toxicus | Bacterial Head Blight |
| Rhizobium rhizogenes | Hairy root disease |
| Rhodococcus fascians | Leafy gall disease |
| Robbsia andropogonis | leaf, bud, and stem spotting |
| Spiroplasma citri | Citrus stubborn disease |
| Spiroplasma kunkelii | Corn stunt disease |
| Strawberry lethal yellows phytoplasma (CPA) | Strawberry lethal yellows disease |

| | |
|---|---|
| Streptomyces acidiscabies | Common Scab, root rot |
| Streptomyces europaeiscabiei | Common Scab |
| Streptomyces scabiei | Common Scab |
| Streptomyces turgidiscabies | Common Scab |
| Trinickia caryophylli | Wilt disease |
| Xanthomonas albilineans | Leaf Scald |
| Xanthomonas arboricola | Necrotic, Leaf Spot |
| Xanthomonas axonopodis | Leaf Spot |
| Xanthomonas campestris | Black rot, Wilt |
| Xanthomonas cassavae | Leaf Spot, necrosis |
| Xanthomonas citri | Citrus canker |
| Xanthomonas cucurbitae | Bacterial Leaf Spot |
| Xanthomonas euvesicatoria | Bacterial Leaf Blight |
| Xanthomonas fragariae | Bacterial Leaf Spot |
| Xanthomonas hortorum | Bacterial Leaf Spot |
| Xanthomonas hyacinthi | Yellow disease |
| Xanthomonas hydrangeae | Bacterial Leaf Spot |
| Xanthomonas oryzae | Blight and leaf streak |
| Xanthomonas perforans | Bacterial Leaf Spot |
| Xanthomonas phaseoli | Leaf Blight |
| Xanthomonas prunicola | Bacterial Leaf Streaks |
| Xanthomonas sacchari | Leaf Chlorotic Streak Disease |
| Xanthomonas theicola | Cankers |
| Xanthomonas translucen | Bacterial Leaf Streak |
| Xanthomonas vasicola | Bacterial Leaf Streak |
| Xanthomonas vesicatoria | Leaf spots, fruit spots and stem cankers |
| Xylella fastidiosa | Bacterial leaf scorch, alfalfa dwarf, chlorosis |
| Xylella taiwanensis | Pear leaf scorch disease |
| Xylophilus ampelinus | Bacterial blight of grapevine, canker of grapevine |
